# Hunting for Helminths: short- and long-read shotgun metagenomics for parasite detection in faecal samples

**DOI:** 10.64898/2026.03.09.710549

**Authors:** Katie O’Brien, Ajay Elamaran, Mehmet Dayi, Georgia Keeling, William D Nevin, Yuchen Liu, Mark Viney, Kieran Reynolds, Charmaine Bishop, Banchob Sripa, Menebere Woubshete, Pamela Sachs Nique, Rhiannon Wright, Jane Younger, Vicky L Hunt

## Abstract

Soil-transmitted helminths (STHs) pose significant challenges to public health in endemic areas, necessitating reliable methods for their detection. Shotgun metagenomics enables simultaneous detection of STHs and microbes in a sample without prior knowledge of what is present. However, validation of shotgun metagenomics with known infection intensity or across different sequencing platforms has not been carried out for eukaryote parasites including STHs, and false positives remain a pervasive issue. We validated shotgun metagenomics as a method of STH detection in faecal samples. Using the *Strongyloides ratti* laboratory model of a STH infection we investigated how analytical methods (nucleotide-nucleotide matching, nucleotide-protein matching, marker gene detection, mitochondrial mapping), infection intensity and sequencing technology (short-read vs. long-read) affects sensitivity and specificity of detection. *S. ratti* was accurately detected at a standard laboratory dose, but low intensity infections were more difficult to detect. Only mitochondrial sequence mapping was 100% accurate at identifying *S. ratti* with no false positives. Overall, short-read outperformed long-read sequencing methods. We applied the same analytical methods to human faecal samples with confirmed infections for at least one of four STHs. Mitochondrial sequence mapping was also the most effective method for detecting STHs in human faecal samples, detecting 100% of *Necator americanus* and 92% of *Ascaris* spp. infections, but could not reliably detect STHs where DNA levels are expected to be low or variable. In conclusion, mitochondrial mapping was the most effective method of detection for sensitivity and specificity in both the laboratory system and human faecal samples. Our findings indicate that shotgun metagenomics should be approached cautiously using validated methods, particularly when infection intensity or DNA levels are expected to be low.

**Author Summary:** Soil-transmitted helminths (STH) such as the parasite *Strongyloides*, are important gastrointestinal parasites of humans and livestock. Accurate methods of detection for diagnostics and monitoring are important to implement suitable control and treatment strategies. Here we validate a shotgun metagenomics approach, where all DNA in a sample is sequenced, for detecting STH in faecal samples using a *Strongyloides* laboratory model for infection. *Strongyloides* was reliability detected in faecal samples at higher infection levels, but mitochondrial genome mapping of the sequences was the only analytical method that reliably detected *Strongyloides* at lower infections levels. These results were reflected in stool samples from humans infected with STH, where mitochondrial mapping was also the most reliable method. However, species that were associated with low levels of parasite material or DNA in the faeces including *Strongyloides stercoralis,* were more difficult to detect. We compared two sequencing methods: short-read Illumina and long-read Oxford Nanopore Technologies, but short-read outperformed long-read shotgun metagenomics. Contamination of bacteria sequences in parasite genome assemblies was problematic for analysis and contributed to false positive results. Future work should focus on specific targeting of eukaryote DNA either at the laboratory or bioinformatic stage to improve STH detection further.

## 1. Introduction

Soil-transmitted helminths (STHs) infect 1.5 billion people globally, particularly in Low- and Middle-Income Countries and are recognised by the WHO as a Neglected Tropical Disease (NTD)^1^. Despite the progress that has been made in reducing both transmission and morbidity caused by STHs through preventative chemotherapy, detection of STH infection remains a challenge.

Coproscopic methods of STH detection^2^ are considered generally reliable^3,4^ and low cost^5^, but they require specialised technicians, are time consuming and low throughput. Where egg and larval outputs vary over time, as is often observed in natural infections, coproscopy can lack sensitivity^6–9^ and underestimate prevalence, particularly for some key species such as the STH *Strongyloides stercoralis*^10^. Molecular techniques such as loop-mediated isothermal amplification (LAMP) assays and PCR/RT-PCR are more sensitive and have begun supplementing conventional STH detection methods^11–15^. However, they are costly, can require *a priori* knowledge of the target species^16^, face challenges such as taxon-specific PCR amplification biases^17,18^ and a lack of resolution for species-level identification^19–21^. STHs commonly occur as co-infections^22^ and symptoms cannot be used to determine the agent of disease^23,24^. An untargeted method of identification is therefore valuable for diagnostics.

Shotgun metagenomics sequences all DNA in a sample, allowing for simultaneous characterisation of eukaryotic, bacterial and viral diversity, without prior knowledge of the organisms which are likely to be present. As such, shotgun methods have the potential to detect multi-STH infections and allow for exploration of the microbiome, including bacterial infections. The use of shotgun metagenomics for detecting microbes is well-established^25–29^, benchmarked, and validated^30–36^. False positive reporting is a serious issue in bacterial pathogen detection, and guidelines to ensure accuracy have been published, highlighting the importance of validating methods^37,38^. Shotgun metagenomics for eukaryote parasite detection lacks validation, and analytical methods are not well established^39–45^. Studies have mostly relied on confirmation of parasite infection by other methods of detection, such as microscopy or targeted PCR, and infection intensity is either unknown or estimated^46–51^. Among the major challenges in using shotgun metagenomics for STH detection are: (i) STHs are present at lower abundance (i.e. smaller fraction of reads in sequencing libraries) than microbes in faecal samples, making them more difficult to detect^52^; (ii) eukaryotic genome assemblies are commonly contaminated with bacterial sequences, which can lead to false positive identifications^53–55^.

The potential of long-read shotgun metagenomic sequencing with Oxford Nanopore Technologies (ONT) or Pacific Biosciences to simultaneously detect and identify eukaryotes and microbes remains largely unexplored^56,57^. These methods offer advantages of portability and real-time data generation (ONT), and could aid detection of STHs that lack high quality genome assemblies by identifying longer sequences. The cost of both long-read and short-read metagenomics is reducing, presenting feasible diagnostic tools for the future^58,59^, as well as use as a monitoring and research tool.

We use a laboratory system of rats infected with known infection intensity of *Strongyloides ratti,* an established laboratory model for *Strongyloides* infection^60^. STH detection using shotgun metagenomics is assessed on the analytical approach used, infection intensity (standard and low dose) and sequencing technology (short-read, long-read). We further apply these methods to human faecal samples infected with one of four STHs and assess the sensitivity and specificity of the methods tested.

## 2. Methods

### Ethics Statement

Rodent work was performed under the authority of licences issued by the Animals (Scientific Procedures) Act 1986 and was approved by the University of Bath Animal Welfare and Ethical Review Body. Collection of samples from human participants was carried out after either written or oral consent. The study was approved by the Khon Kaen University Ethics Committee for Human Research (reference HE641605) and the Central University Research Ethics Committee at the University of Liverpool (reference 10936).

### Maintenance of Strongyloides ratti

Five week old female Wistar rats were infected with *S. ratti* ED321 by subcutaneous injection of third stage infective larvae (iL3) prepared from faecal cultures as described by Hunt et *al*.^61^ Rats were infected with either 50 (low dose) or 500 (standard dose) iL3s suspended in 200ul of PBS. Controls were injected with PBS only.

### Sample collection and preparation

i. ***S. ratti***: Rat faeces was collected from rats between 6 - 11 days post infection (dpi) and stored at -80°C until extraction. DNA extraction was carried out using the QIAamp PowerFecal Pro DNA kit (Qiagen) according to the manufacturer’s instructions (**Additional File 1**).
ii. ***S. stercoralis*** :Four human stool samples (CKH130, CKH21, LWH76, KKU01) were collected from participants in Thailand (**Table 1**) as described by Viney et *al*.^62^

**Table 1:** *Strongyloides stercoralis*-positive samples collected from Thailand.

| Sample ID | Sample Site | Age | Gender | No. iL3s |
| --- | --- | --- | --- | --- |
| CKH21 | 16°07'53"N 102°39'01"E | 71 | Male | >1000 |
| LWH76 | 16°07'53"N 102°39'01"E | 62 | Male | >500 |
| CKH130 | 16°07'53"N 102°39'01"E | 50 | Male | 45 |
| KKU01 | 16°28'11"N 102°49'52"E | Unknown | Unknown | 53 |

Three further samples were collected from patients in the UK who had recently migrated from Fiji (infective larvae numbers present in faeces is unknown for these samples). All human participants were confirmed to have a *S. stercoralis* infection using microscopy or PCR (**Additional File 1**).

### Illumina short-read sequencing

DNA fragment libraries from standard dose rat faeces were constructed with NEBNext Ultra II FS DNA kit (1/2 volume protocol). DNA fragment libraries from low dose infection rat faeces and human faecal samples (Thailand and Fiji) were constructed by Eurofins Genomics. Sequencing was carried out on a NovaSeq SP (2×150bp), generating ∼10-12.9 million (rat) and ∼41.5 million (human) paired end reads per library (**Supplementary Table S1-S4**, **Additional File 1**). Host genome sequences (*Rattus norvegicus* Rnor 6.0, NIH accession GCF_000001895.5) were removed using Bowtie2 (v. 2.4.2)^63^ and Samtools (v. 1.1.2)^64^. The Bedtools (v. 2.29.2)^65^ ‘bamtofastq’ was used to convert bam files into fastq.

### ONT sequencing and pre-processing

A subset of three samples (one infected 11dpi standard dose, one infected 11dpi low dose and one control 11dpi) used for short-read sequencing were selected for long-read metagenomic sequencing using MinION sequencing technology from Oxford Nanopore Technologies (ONT). Samples were prepared for sequencing using the SQK-LSK110 Ligation Sequencing Kit according to the manufacturer’s instructions in combination with the following NEBNext products: NEBNext FFPE Repair Mix (M6630), NEBNext Ultra II End repair/dA-tailing Module (E7546) and the NEBNext Quick Ligation Module (E6056). Samples were sequenced for up to 72 hours on a FLO-MIN106 flow cell connected to a laptop hosting MinKNOW software. The fast5 files generated from sequencing were converted into fastq format using the high accuracy version of Oxford Nanopore’s GPU-enabled ‘Guppy’ program with the following parameters: flowcell:‘FLO-MIN106’, kit:‘SQK-LSK110’, callers:14, gpu runners per device:8, chunks per runner:768, chunk size:500. Quality control assessment of the MinION run was performed using MinIONQC^66^ and FASTQC^67^, and the first 35 bases of each sequencing read were removed due to low quality using NanoFilt^68^. All reads greater than 54,000 bases in length were also removed due to low quality. Following trimming, reads were processed by Nanoplot^68^, and sequences => Q7 were retained for further analysis. Host genome sequences (from same source as short-read analysis) were removed using Minimap2 with the ‘ax map-ont’ preset^69^ and the resulting unmapped reads were converted into fastq files as previously mentioned (**Supplementary Table S5, Additional File 1**).

### Detection of *S. ratti* using short-read metagenomics

Taxonomic assignment was compared using four analytical tools: Kraken2 (nucleotide-nucleotide matching)^70^, DIAMOND+MEGAN (nucleotide-protein matching)^71^, EukDetect (marker genes)^72^, and mitochondrial mapping^47^ (**Additional File 1 & 2)**.

### i. K-mer based nucleotide-nucleotide matching with Kraken2

A custom Kraken2 nucleotide reference database was constructed using the ‘kraken2-build’ command. The standard RefSeq libraries supplied by Kraken2 for fungi, archaea, bacteria, viruses, and protozoa were first downloaded using the ‘kraken2-build --download-library’ command. Following this, the reference sequences for the host (*Rattus norvegicus* - Rnor6.0) and its main dietary components (wheat: IWGSC CS RefSeq v2.1, barley: MorexV3_pseudomolecules_assembly, soy: Glycine_max_v4.0 and maize: Zm-B73-REFERENCE-NAM-5.0) were downloaded from NCBI and added to the database using the ‘kraken2-build --add-to-library’ command. The genomes for *S. ratti* and five closely related species, alongside a further eight nematode genomes that span multiple evolutionary clades of Nematoda, were also added to the database. The nematode genomes included were: *S. ratti, S. stercoralis, Parastrongyloides trichosuri, S. papillosus, S. venezuelensis, Rhabditophanes sp. KR3021, Meloidogyne hapla, Bursaphelenchus xylophilus, Ascaris suum, Brugia malayi, Trichuris muris, Trichinella spiralis, Necator americanus* and *Caenorhabditis elegans* (**Supplementary Table S6, Additional File 1**). The addition of these genomes enabled the assessment of false positive hits to non-target genomes. All nematode genomes were downloaded from the WormBase parasite database V16.0^73^ (accessed 13/05/2022). The final custom database was constructed using the ‘kraken2-build --build’ command.

Kraken2 classification was applied to all short reads against the custom database, using the ‘--report’, ‘--classified-out’, and ‘--gzip-compressed’ options. All Kraken2 ‘report’ output files were combined into a biom file alongside the relevant metadata using kraken-biom (v. 1.0.1). The biom file was imported into R (v. 4.2.1) using the ‘import-biom’ function from phyloseq v.1.46.0^74^.

A custom R script was run to prevent erroneous taxonomic assignments due to taxonomic revisions (https://github.com/Vicky-Hunt-Lab/Metagenomics_tax_update). This involved taking the taxa IDs from the OTU table and reclassifying them according to current NCBI taxonomy using the ‘classification’ function from the rentrez package (v.4.3.2)^75^ in R, e.g. all entries classified as phylum “Firmicutes” were reclassified to “Bacillota”. An abundance filter was applied to remove any observations reaching less than 0.001% of total abundance. A variety of alpha diversity measures were calculated to assess within-sample diversity between infected and control samples, using the ‘estimate_richness’ function from phyloseq. Comparisons of alpha diversity between control and infected sample groups were performed using ANOVA .

Beta diversity measures were estimated using the ‘ordinate’ function in phyloseq, employing principal coordinate analysis (PCoA) to visualise pairwise Bray-Curtis distances. Permutational multivariate analysis of variance (PERMANOVA) tests, conducted using the ‘adonis2’ function in Vegan (v. 2.6.2)^76^, were used to evaluate dissimilarities in community composition between infected and control samples. Taxonomic composition was visualised using ggplot2 (v.3.3.6)^77^ in R. To investigate the rate of true positive assignment of nematode taxa using Kraken2, any hits to the phylum “Nematoda” were filtered, and reads not aligning to a specific species were discarded.

### ii. Nucleotide-protein matching with DIAMOND+MEGAN pipeline

We followed the methods outlined in Bağcı et *al*. ^71^ where short-read sequencing outputs were translated, then aligned against the NCBI-nr database using the ‘blastx’ function in DIAMOND (v. 2.0.15)^78^, followed by taxonomic binning with MEGAN community edition (v. 6.24.11)^71^. DIAMOND’s double-indexing approach is approximately 20,000 times faster than a regular blastx search^78^. The nr database was downloaded from NCBI (12^th^ May 2022), and a DIAMOND index was created using the ‘diamond makedb’ command. After the concatenation of paired-end samples, the ‘diamond blastx’ command was used to produce a diamond archive file (.daa). The MEGAN database (megan-map-Feb2022.db) and tools suite were downloaded from the MEGAN website^79^. The ‘daa-meganizer’ function categorised reads into taxonomic bins. Alignments were converted for analysis using the pb-metagenomics-tools script ‘Convert_MEGAN-NCBI-c2c_to_kreport-mpa.py’ (https://github.com/PacificBiosciences/pb-metagenomics-tools). The “megan-c2c.kreport.txt” output files were combined into a biom file alongside the relevant metadata using kraken-biom (v. 1.0.1) and was imported into R as described previously.

Eukaryotic reads were extracted from the .daa files using the ‘read-extractor’ function from MEGAN, followed by rerunning the pipeline with the ‘--iterate’ option for increased sensitivity in the blastx search. The minimum support percentage for taxonomic assignment after eukaryotic extraction was adjusted from the default of 0.01% to 0.1% to provide more conservative assignments. All outputs from the eukaryotic extraction step were again combined into a biom file, and any species-level alignments to taxa falling under the phylum “Nematoda” were retained and imported into R as described previously. The DIAMOND+MEGAN pipeline was not feasible for high depth sequencing analysis due to computational demands (**Additional File 1**). As such, we do not recommend using this method for high depth sequencing where samples generate more than 50 million reads.

### iii. Eukaryote marker gene detection with EukDetect

To investigate whether species-specific markers could be used in the identification of *S. ratti*, EukDetect (v. 2, 14^th^ October 2022) was used^72^. Of the >500,000 markers included in the database, approximately 17,000 are derived from >150 species of nematode or platyhelminth, including 13 of the 14 nematodes chosen for inclusion in the custom Kraken2 database (*A. suum* is not included). The EukDetect database was downloaded from Figshare (v. 9)^80^ and the analysis was conducted using the ‘runall’ mode on all short reads.

### iv. Mitochondrial mapping

In our Kraken reference database, only three nematode species - *S. stercoralis*, *T. muris*, and *T. trichiura* - had mitochondrial genomes included with their nuclear reference assemblies when downloaded from their repository source. To evaluate whether mitochondrial mapping is a valid detection approach, we therefore performed a separate analysis. Mitochondrial mapping was performed as per Papaiakovou et *al*.^47^, with the exception that Bowtie2^63^, rather than BWA-MEM^81^, was used for mapping reads. Analysis in R followed the script https://github.com/MarinaSci/IJP_genome_skimming. Alignments to genomes are reported as “normalised coverage” which refers to the number of reads mapped per million reads generated per mitochondrial genome size (Mb).

A normalised “reads aligned per million reads generated” value was created for all species-level nematode alignments (RPM), to adjust for variation in read counts across samples for all Kraken2, DIAMOND+MEGAN and EukDetect outputs, alongside the “Normalised coverage” values generated using mitochondrial mapping (**Additional File 2**). Heatmap visualisation for all four bioinformatic methods were generated using ggplot2 in R. Wilcoxon rank sum tests were conducted to compare the RPM of alignments between various sample groups.

Note: A comparison of computational effort between all four methods applied here can be found in **Additional File 1**.

### Taxonomic assignment and detection of *S. ratti* using long-read shotgun metagenomic reads

We used the following tools for long-read data analysis: i) nucleotide-protein matching with DIAMOND+MEGAN-LR^71^, ii) Assembly-based alignment with the CZ ID online platform^82^, and iii) mitochondrial mapping.

### i. Nucleotide-protein matching with DIAMOND+MEGAN pipeline

The DIAMOND+MEGAN pipeline was used for long-read analysis with alterations to account for longer reads. Firstly, we applied a recommended penalty for frameshifts in DNA-vs-protein alignments (‘-F 15’). We also added the parameters ‘--range-culling’ and ‘--top 10’, enabling the filtering of alignments below a certain bit score, thus retaining only the most relevant hits across a query sequence. Diamond archive output files were meganized using ‘daa-meganizer’ with the ‘--longReads’ parameter so that the number of aligned bases, rather than reads, would be used to assign taxonomy. As a result, all taxonomic assignments for long-read data are reported as “bases per million” (BPM). An abundance filter was applied to remove any observations reaching less than 0.001% of total abundance. Eukaryote read extraction, and conversion into biom files remains the same as for short reads.

### ii. Assembly-based alignment with CZ ID online platform

Long-read sequences were uploaded to CZ ID^83^, where they were profiled using the mNGS nanopore pipeline. Species-level taxonomic assignments were imported into R and taxonomic categories were reformatted as described previously (https://github.com/Vicky-Hunt-Lab/Metagenomics_tax_update) and BPM values were generated (**Additional File 3**).

### iii. Mitochondrial mapping

Mitochondrial mapping was performed as per Papaiakovou et *al* ^47^. For long-read samples, both Bowtie2 and Minimap2 ^69^ were used to map reads, to assess performance using a short-read aligner (Bowtie2) compared to a long-read aligner (Minimap2). Analysis in R followed the script in https://github.com/MarinaSci/IJP_genome_skimming. Alignments to genomes are reported as “normalised coverage” which refers to the number of reads mapped per million reads generated per mitochondrial genome size (Mb) (**Additional File 3**).

### Detection of *S. ratti* using hybrid metagenomic assemblies

For the three samples that had both short- and long-read sequencing data (Standard Dose 11dpi, Low dose 11dpi, Control 11dpi), hybrid assemblies were generated using metaSPades v3.15.5 with default options^84^. Assemblies were further evaluated by metaquast v5.2.0 with default options^85^ and contigs were grouped into bins using MetaBAT2 v2.12.1. with default options^86^ where each bin corresponds to an individual genome. Assembly bins were evaluated for contamination, completeness and N50 values with CheckM v1.2.2^87^, and those with contamination <10% and completeness >90% were considered complete genomes (**Supplementary Table S7, Additional File 1**). Three files were generated from hybrid assembly (i) a file containing all assembled contigs including unassembled reads, (ii) a MetaBAT2 binned file containing bins clustered into putative genomes, (iii) a MetaBAT2 unbinned sequence file containing all sequences not assigned to a genome bin. The assembled contigs for each sample were aligned against a custom Kraken2 database as mentioned previously. Nematode classifications were normalised to RPM (**Additional File 3**). Mitochondrial mapping was also performed as mentioned previously for long reads, using both Bowtie2 and Minimap2, to generate normalised coverage values (**Additional File 3**).

A custom BLAST database was created using the “makeblastdb” command with default parameters comprising the Rnor 6.0 genome, the *S. ratti* genome, and 13 other nematode species from four evolutionary clades including free-living and parasitic nematodes, as included in the Kraken2 custom database. As all binned sequences had microbial origins (**Additional File 4**), the BLAST search was performed on unbinned sequences with the ‘blastn’ command, using an e-value threshold of 0.001 and a percent identity of 95% (**Additional File 5**). In the first instance, the BLAST search was carried out only on sequences greater than 1kb in length. Subsequently, another BLAST search was also performed on sequences less than 1kb in length to do a comparative analysis.

### Sensitivity, specificity rates

Measures of sensitivity, specificity, precision, false positive detection rate (FPR), and F1 scores for short-read (Additional File 6), long-read and long-short read hybrid analysis (Additional File 7) were calculated as below:

Sensitivity = True Positives/(True Positives + False Negatives)

Specificity = True Negatives/(False Positives + True Negatives)

Precision = True Positives/(True Positives + False Positives)

FPR = 1 – Specificity

F1 Score = 2 * (Sensitivity* Precision) / (Sensitivity + Precision).

### Parasite detection using short-read metagenomics on human samples

We reanalysed 24 metagenomic shotgun samples (SRA, BioProject PRJNA407815) from a previously published microbiota-parasite interaction study^88^. Kraken2, EukDetect, and mitochondrial mapping were applied as described above for short-read analysis of *S. ratti* in rat faecal samples (**Additional File 8**). DIAMOND+MEGAN was not used because it was the least accurate for *S. ratti* samples (**Additional File 6**) and computationally more demanding (**Additional File 1**). Results were compared with qPCR or formal-ether results (**Supplementary Table S8, Additional File 1**) for *Necator americanus*, *Ascaris spp*., and *Trichuris trichiura* infections. True-positive RPM detection rates, using sensitivity, specificity, precision, false positive detection rate, and F1 scores, were calculated as described above (**Additional File 9**).

The following adjustments were made prior to analysis of human faecal samples: 1) trimmomatic v. 0.39^89^ was used to remove any unpaired or short (<20bp) reads, 2) An alignment length filter of 90bp, rather than the default 130bp, was used for mitochondrial mapping, as the files uploaded to SRA had already undergone trimming, 3) The genomes for *Ascaris lumbricoides* and *Trichuris trichiura* were added to the custom Kraken2 database, 4) Any reference genome sequences with length <10 kb were filtered from the Kraken2 reference database. We did this as per Papaiakovou et *al.*^47^, where removal of short scaffolds (<10kb) was shown to effectively remove bacterial contamination from the reference genome of *E. vermicularis* and 5) A minimum threshold of 50 RPM was applied to Kraken2 outputs based on our short-read analysis in controlled lab settings. The *A. lumbricoides* genome was downloaded from WormBase Parasite V16.0 and the *T. trichiura* genome was sourced fromDoyle et *al.*^90^.

We extended our analyses to include the detection of *Strongyloides stercoralis* in seven confirmed positive samples collected from humans in Thailand (n=4) and from participants of a clinical cross-sectional study on Fijian migrants to the United Kingdom (n=3). Kraken2, EukDetect, and mitochondrial mapping methods were used to assign the presence of *S. stercoralis*, using the same custom Kraken2 reference database as in the previous section (with a threshold of 50 RPM and removal of <10kb scaffolds).

## 3. Results

We assessed the importance of the following parameters for detection of *S. ratti* DNA in rat host faecal samples using shotgun metagenomics.

i. Computational analysis: four approaches of taxonomic assignment were used to identify *S. ratti* DNA: 1) Kraken2, a k-mer based nucleotide-nucleotide matching tool^70^, 2) MEGAN+DIAMOND, a nucleotide-protein matching tool^71^, 3) EukDetect, a eukaryote marker gene detection tool^72^, and 4) mitochondrial genome mapping^47^.
ii. Infection intensity: a standard laboratory dose (500 iL3) compared with a low dose (50 iL3) infection.
iii. Sequencing technology: Short-read shotgun metagenomics compared with ONT long-read shotgun metagenomics and hybrid assembly.

### 2.1 *S. ratti* is detected at standard infection dose using shotgun metagenomics

Measurements of alpha and beta diversity showed no significant differences in overall diversity or taxonomic composition between infected and uninfected control samples (**Supplementary Figures S1-S2, Additional File 1**). The majority of reads were classified as ‘bacteria’ and only 0.35% and 0.064% of reads were identified as ‘nematode’ using Kraken2^70^ (**Fig. 1a**) and DIAMOND+MEGAN^71^, respectively (**Fig. 1b**). Kraken2^70^, DIAMOND+MEGAN^71^, EukDetect^72^ and mitochondrial genome mapping^47^ all accurately identified *S. ratti* in the shotgun metagenomic sequencing data from faeces of rats infected with a standard laboratory dose. False positive assignment of reads occurred at low levels using Kraken2 and DIAMOND+MEGAN, but this was resolved by implementing a threshold to remove sequences with <50 reads per million (RPM). All methods had high specificity and sensitivity for detecting *S. ratti* and did not detect false positives (**Fig. 1c-f, Additional Files 2,6**).

**Figure 1.**
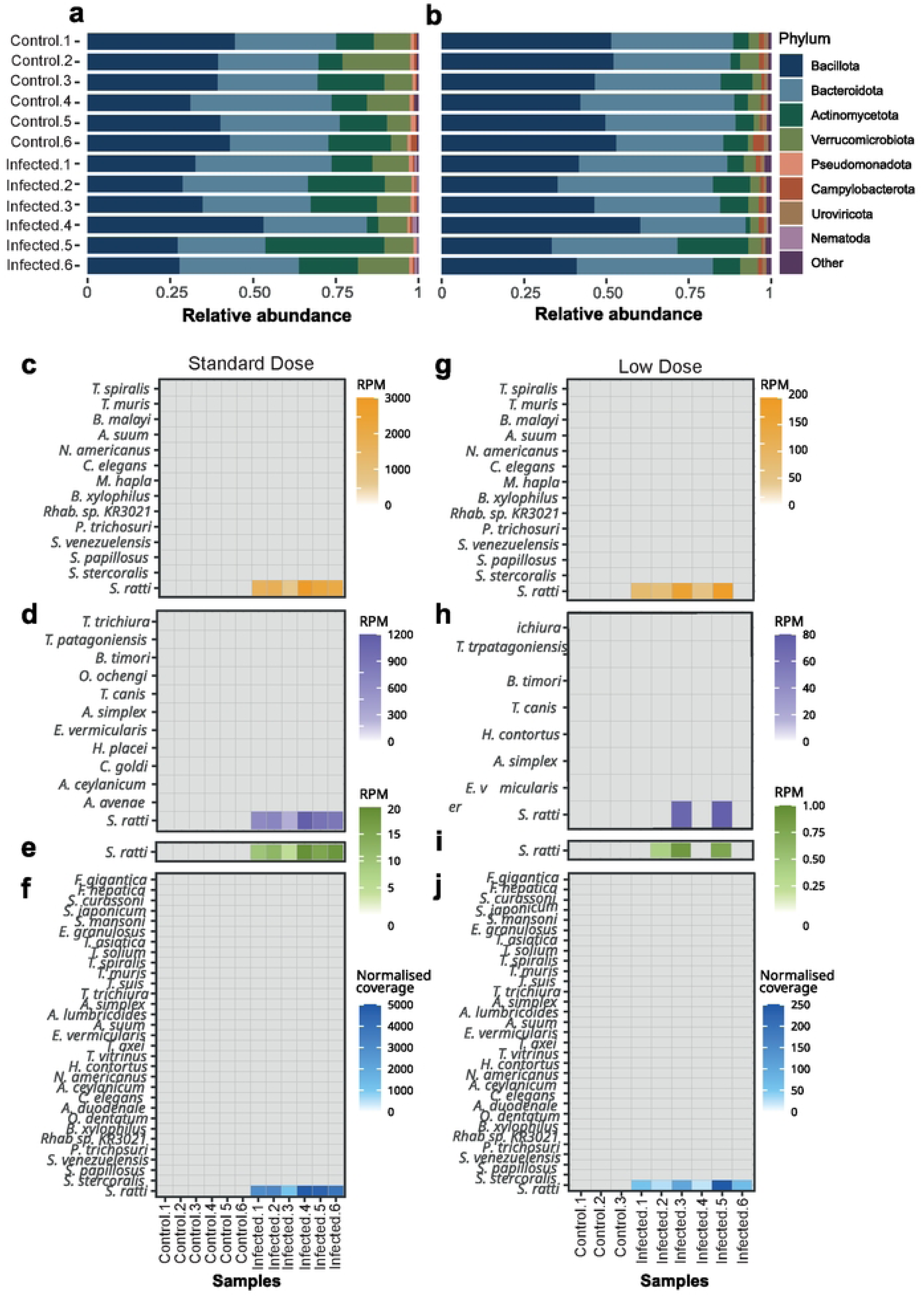
Taxonomic classification and *S. ratti* detection in rat faecal samples using short-read shotgun metagenomics. (**a-b**) Relative abundances of core phyla (>0.001% abundance) in rat faecal samples infected with a standard dose of *Strongyloides ratti* (500 iL3s) with an average sequencing coverage of ∼10.5M reads, identified using (**a**) a custom Kraken2 database and (**b**) the DIAMOND+MEGAN pipeline. Low abundance phyla are grouped into the “Other” category. (**c-f**) assignment of reads to nematodes from short-read shotgun metagenomics rat faecal samples from a standard dose infection of *S. ratti* using (**c**) a custom Kraken2 database containing 14 nematode genomes with a 50 RPM threshold for detection, (**d**) the DIAMOND+MEGAN pipeline with a 50 RPM threshold for detection, (**e**) EukDetect marker gene detection and (**f**) mapping against 31 nematode mitochondrial genomes. (**g-j**) Assignment of reads to nematodes from short-read shotgun metagenomics rat faecal samples from a low dose infection of *S. ratti* using (**g**) a custom Kraken2 database containing 14 nematode genomes with a 50 RPM threshold for detection, (**h**) the DIAMOND+MEGAN pipeline with a 50 RPM threshold for detection, (**i**) EukDetect marker gene detection and (**j**) mapping against 31 nematode mitochondrial genomes. Species are ordered by increasing relatedness to *S. ratti*. Colour on the heatmap denotes level of detection, with grey squares indicating no detection. RPM = Reads mapped per million reads; Normalised coverage = Reads mapped per million reads per genome size (Mb).

### 2.2 *S. ratti* can be detected at low infection intensity using mitochondrial mapping

To test if infection intensity is important for STH detection we compared shotgun metagenomics data from rat faecal samples for a standard dose (500 iL3) to a low dose (50 iL3) of *S. ratti* infection. *S. ratti* was detected at lower levels in samples with lower infection intensity (Wilcoxon rank sum test, W=36, p<0.01) (**Supplementary Table S9**).

Kraken2, DIAMOND+MEGAN and EukDetect produced false negatives, detecting *S. ratti* in 5/6 (**Fig. 1g**, F1=0.91, FPR=0%), 2/6 (**Fig. 1h**, F1=0.5, FPR=0%) and 3/6 (**Fig. 1i**, F1=0.67, FPR=0%) low dose infection samples, respectively (**Additional File 2,6**). As with standard dose samples, a 50 RPM filter was implemented for Kraken2 and DIAMOND+MEGAN to reduce false positive assignment of reads. Reducing the filter threshold to 25 RPM for DIAMOND+MEGAN increased true positive detection of *S. ratti* (F1=0.67), but also increased false positive detection of non-target species (**Additional File 6**). *S. ratti* was correctly detected in all low dose infected samples using mitochondrial mapping and there was no false positive detection of non-target species or detection of *S. ratti* in the control samples (**Fig. 1j**, F1=1, FPR=0%, **Additional File 6**). Overall, low doses of infection lead to poorer STH detection, except for mitochondrial mapping methods which maintained sensitivity and specificity.

### 2.3 Short-read metagenomics outperforms long-read metagenomics and hybrid assembly for detection of *S. ratti*

We assessed the detection of *S. ratti* using ONT long-read metagenomics with two approaches: (i) long-read only (**Fig. 2a-f**), and (ii) long- and short-read hybrid assembly (**Fig. 2g-j**). The same DNA samples were directly compared for long-read and short-read sequencing, so differences are due to the sequencing or analytical methods, not variation between samples.

**Figure 2.**
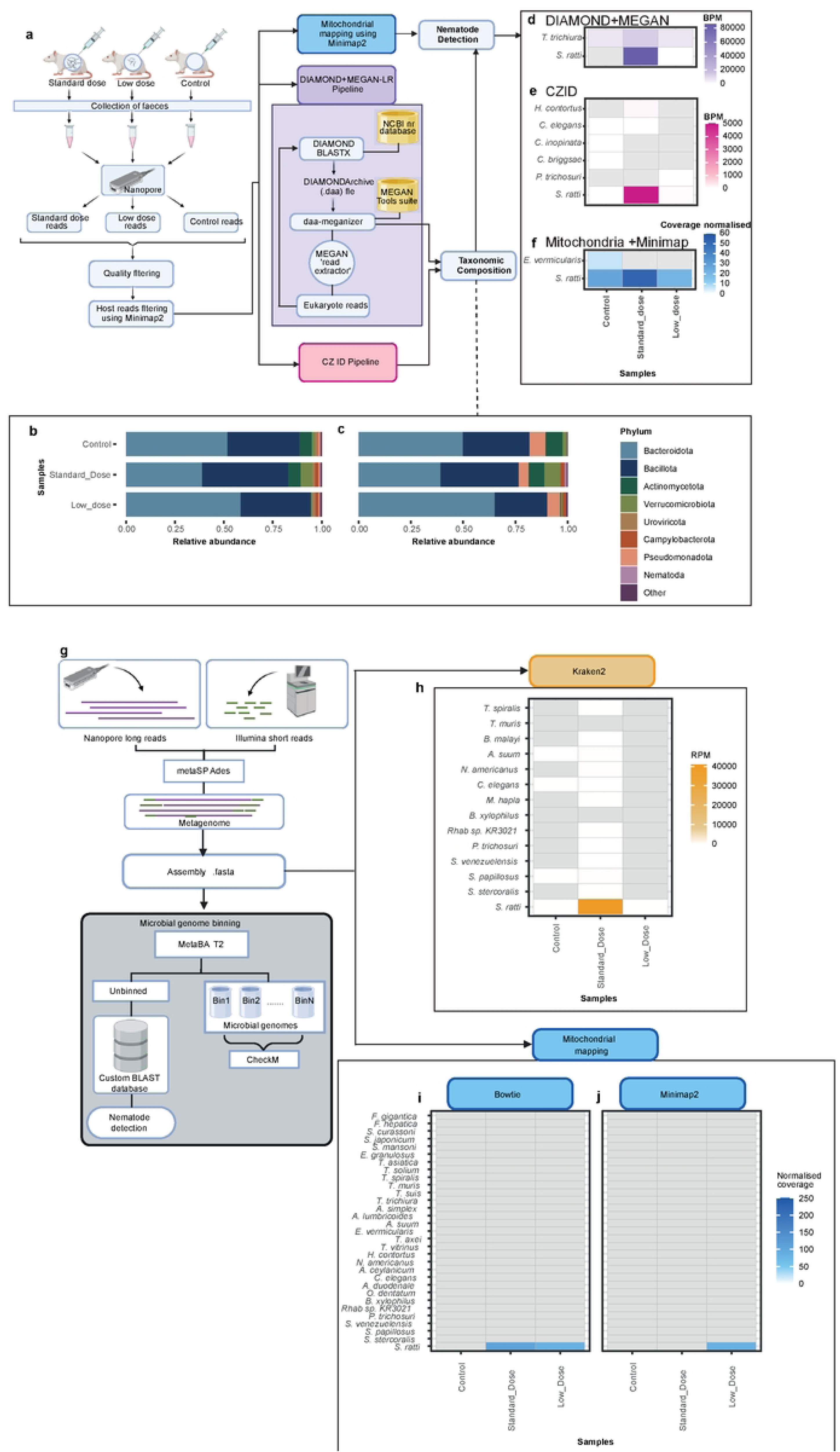
*S. ratti* Detection in Rat Faecal Samples Using Long-Read Shotgun Metagenomics and Long and Short-read Hybrid Metagenomic Assembly. (**a**) Overview of the bioinformatics pipeline used for nematode detection in long-read shotgun metagenomics samples. Three samples were taken at 11 days post infection (11dpi). Sequencing reads were first subjected to trimming and quality filtering and host read removal before analysis using mitochondrial mapping, the DIAMOND+MEGAN-LR pipeline and the CZ ID pipeline. (**b-c**) Relative abundances of core phyla (>0.001% abundance) in rat faecal samples infected with a standard (500 iL3s) or low dose (50 iL3s) of *S. ratti* using ONT long-read sequencing, identified using (**b**) the DIAMOND+MEGAN-LR pipeline and (**c**) the CZ ID pipeline. Low abundance phyla are grouped into the “Other” category. (**d-f**) Nematode detection in long-read shotgun metagenomics samples using (**d**) the DIAMOND+MEGAN-LR pipeline, (**e**) the CZ ID pipeline, and (**f**) mitochondrial mapping with Minimap2. BPM = Bases mapped per million reads; Normalised coverage = Reads mapped per million reads per genome size (Mb). Colour denotes level of detection, with grey squares indicating no detection. (**g**) Overview of the bioinformatics pipeline used for nematode detection using hybrid assemblies of short- and long-read shotgun metagenomics. Four methods of detection were employed, using a custom BLAST database against unbinned assembled contigs, Kraken2, and mitochondrial mapping using Bowtie2 and Minimap2. (**h-j**) Nematode detection in hybrid metagenomic assemblies infected with a standard (500 iL3s) or low dose (50 iL3s) of *Strongyloides ratti*, using (**h**) a custom Kraken2 database containing 14 nematode genomes, (**i**) mapping against 31 nematode mitochondrial genomes with Bowtie2 and (**j**) Minimap2. RPM = Reads per million reads; Normalised coverage = Reads mapped per million reads per genome size (Mb). Colour denotes level of detection, with grey squares indicating no detection.

The long-read data identified 0.1% and 0.3% ‘nematode’ reads for DIAMOND + MEGAN and CZ ID, respectively for a standard dose infection (**Fig. 2b,c**). DIAMOND+MEGAN and CZ ID detected *S. ratti* at the standard and low dose (**Fig. 2d,e**). However, false positive assignments to *T. trichiura* were observed in all long-read samples using DIAMOND+MEGAN (**Fig. 2 d**). Five species were falsely identified by CZ ID and *S. ratti* was detected in control samples (**Fig. 2e**). Applying a BPM threshold to eliminate false positives also lead to false negatives. Mitochondrial mapping with Bowtie2 produced no alignments to *S. ratti* in standard or low dose infections. Mitochondrial mapping with Minimap2 was also unsuccessful (F1=0), because the control sample falsely detected *S. ratti* in control samples and *Enterobius vermicularis* (**Fig. 2f**) (**Additional File 3,7,5**).

A short-long-read hybrid assembly generated at least 445,183 contigs per sample and a total assembly size of >435 Mb/sample (**Supplementary Table S7, Additional File 1**). As no standard method for assignment of assembled contigs has been established for eukaryotic sequences, we compared several approaches (**Fig. 2g**):

i. Kraken2 kmer approach: *S. ratti* was identified at 40,505 RPM for the standard infection dose (**Fig. 2h**), and only 61 RPM for the low infection dose, a similar level to the false positive identification of *S. ratti* in the control sample (74 RPM), suggesting that this method is not suitable for detecting low intensity *S. ratti* infection.
ii. Mitochondrial mapping: *S. ratti* was correctly identified using Bowtie2 (normalised coverage value standard = 109, low = 86.1). *S. ratti* was only identified in the low dose sample using Minimap2 (normalised coverage value=86.1). Neither Minimap2 nor Bowtie2 mapping methods produced false positive alignments. Hybrid mitochondrial mapping outperformed long-read only mitochondrial mapping (Hybrid: F1=1, Long: F1=0) (**Fig. 2i,j**).
iii. Genome assignment of contigs: Contigs were assigned to genomes (METABAT2, CheckM) and unassigned contigs were identified using BLAST: Assembled contigs were assigned to 291 (standard dose), 165 (low dose) and 143 (control) genomes, including 47 (standard dose), 15 (low dose) and 15 (control) full genome assemblies, and were all taxonomically assigned as microbes (**Additional File 4**) (**Supplementary Table S7, Additional File 1**). The standard dose sample had only 46 hits to *S. ratti* for contigs >1kb in length, with alignments between 0.9 to 5.7kb (**Additional File 5**). No hits were identified for *S. ratti* in the low dose and control sample. Only hits for *S. ratti* and the host organism *Rattus norvegicus* were identified across samples (FPR=0). Expanding our criteria to include sequences <1kb identified 15,144 and 30 *S. ratti* hits for standard and low dose samples, respectively. No *S. ratti* hits were identified in the control sample, but false positive detection of non-target nematodes were detected at low levels in all samples (**Additional File 5**). This method lacked sensitivity at low doses. Overall, long-read ONT sequencing offered no advantage for detecting *S. ratti* compared with short read shotgun metagenomics.

### 2.4 Shotgun metagenomics identifies *Necator americanus* and *Ascaris spp.* in human faecal samples

We analysed previously published sequence data for 24 shotgun metagenomic samples of human faeces collected from Indonesia and Liberia^88^ using EukDetect, Kraken2 (50RPM threshold) and mitochondrial mapping. Of the 24 samples, 15, 12 and 2 were qPCR-positive or formol ether concentration positive for *N. americanus, A. lumbricoides* and *T. trichiura*, respectively^88^ (**Supplementary Table S8, Additional File 1**).

Of the 15 confirmed *N. americanus* samples, Kraken2, EukDetect, and mitochondrial mapping methods correctly identified *N. americanus* in 11, 6 and 15 samples, respectively (**Fig. 3a**). False positive assignment to *N. americanus* using Kraken2 occurred in 3 out of 24 samples (**Fig. 3a,b**) (F1 score = 0.76, FPR=33%). Mitochondrial mapping and EukDetect only detected *N. americanus* in qPCR confirmed samples (FPR=0%), and produced F1 scores of 1, and 0.57 respectively. For low-intensity infections (based on qPCR), EukDetect performed the worst, failing to detect all three low-level samples; Kraken2 missed one and mitochondrial mapping missed no low-intensity infections (**Fig. 3a-d; Additional File 8,9**).

**Figure 3.**
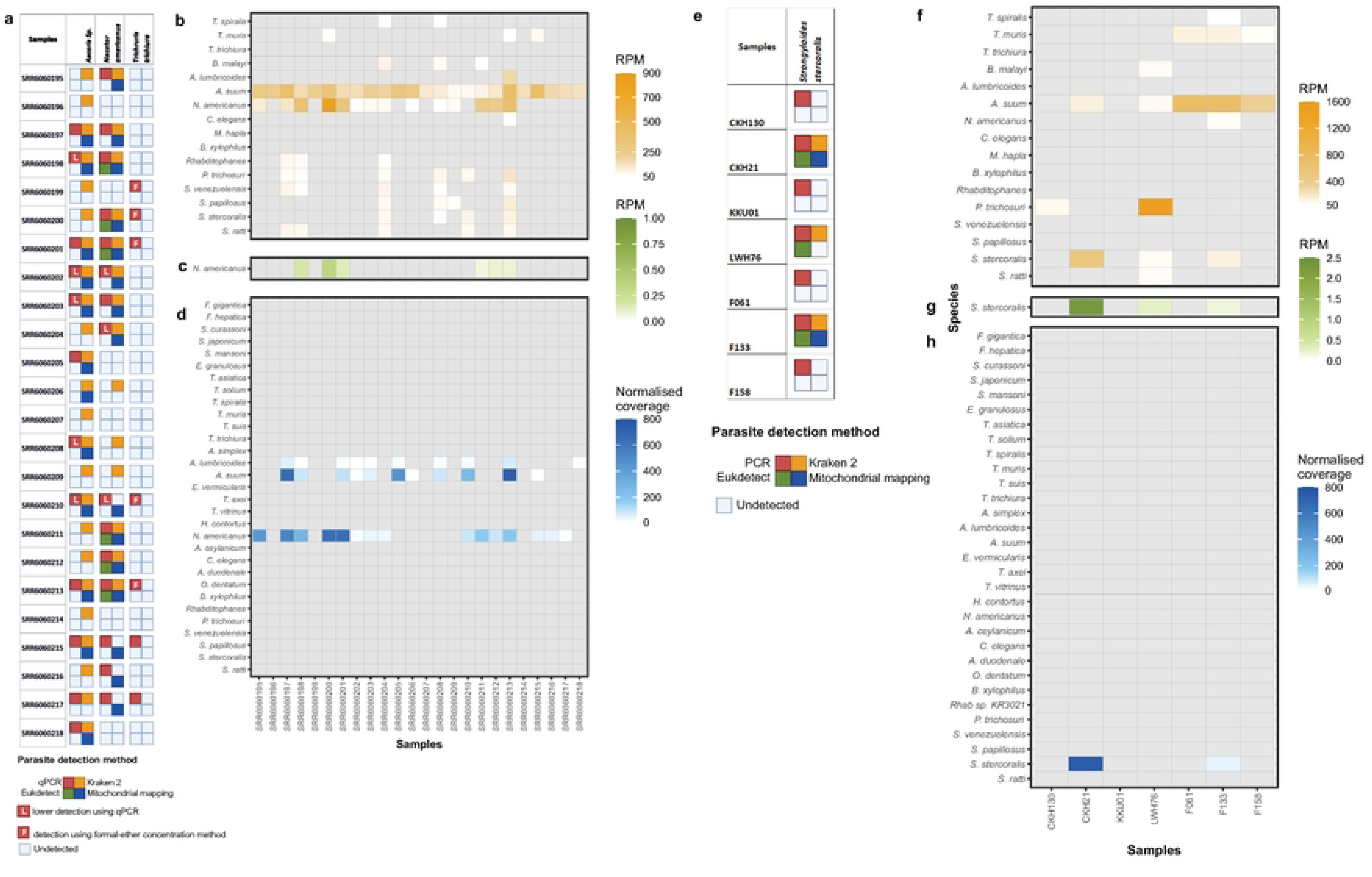
STH Detection in Human Faecal Samples. (**a**) Comparison of qPCR/formal ether concentration assays and three bioinformatics methods used on shotgun metagenomic sequences for the detection of *Ascaris spp., Necator americanus,* and *Trichuris trichiura* in human faecal samples. Red squares indicate confirmed infection using qPCR/formal-ether concentration methods, “L” refers to samples designated as having “low” levels of detection in qPCR, “F” refers to detection using formal-ether concentration. (**b-d**) Nematode detection in short-read shotgun metagenomics samples sourced from infected and uninfected human faecal samples with *Ascaris spp.*, *N. americanus* and/or *T. trichiura* using (**b**) a custom Kraken2 database containing 16 nematode genomes including the three target species with a 50 RPM threshold for detection and scaffolds <10kb removed (**c**) EukDetect marker gene detection and (**d**) mapping against 31 nematode mitochondrial genomes. RPM = Reads mapped per million reads; Normalised coverage = Reads mapped per million reads per genome size (Mb). Colour denotes level of detection, with grey squares indicating no detection. Note: *Ascaris spp.* are not included in the EukDetect database. (**e**) Comparison of qPCR/microscopy assays and three bioinformatics methods used on shotgun metagenomic sequences for the detection of *S. stercoralis* in human faecal samples. Red squares indicated confirmed infection. Samples CKH130, CKH21, KKU01 and LWH76 were collected from in Thailand, samples F061, F133 and F158 were collected from people travelling to the UK from Fiji. (**f-h**) Nematode detection in short-read shotgun metagenomics samples sourced from individuals infected with *S. stercoralis* using (**f**) a custom Kraken2 database containing 16 nematode species, with a 50 RPM threshold for detection and scaffolds <10kb removed from consideration, (**g**) EukDetect marker gene detection and (**h**) mapping against 31 nematode mitochondrial genomes. RPM = Reads mapped per million reads; Normalised coverage = Reads mapped per million reads per genome size (Mb). Colour denotes level of detection, with grey squares indicating no detection.

*Ascaris spp.* was detected in all qPCR-positive and -negative samples using Kraken2 (F1=0.66 , FPR=100%) (**Fig. 3a,b**). Mitochondrial mapping detected *Ascaris spp.* in 11 of the 12 qPCR positive samples and produced fewer false positive assignments (**Fig. 3a,d**, FPR=0.08%), providing more reliable *Ascaris spp.* detection (F1=0.91). Kraken2 and mitochondrial mapping successfully detected all low-level infections. Read assignments were more common for *A. suum* than *A. lumbricoides*, however, qPCR confirmation was only tested for *A. lumbricoides. Ascaris* spp. was therefore considered here at the genus level. *T. trichiura* was the only species that was not detected by either Kraken2, EukDetect, or mitochondrial genome mapping (**Fig. 3a-d**). Together our results support that mitochondrial mapping is the most accurate and reliable approach to identify *N. americanus* and *Ascaris spp.* infections. EukDetect had the most false negative results and Kraken2 had the most false positives (**Additional File 9**).

### 2.5 Shotgun metagenomics did not reliably detect *S. stercoralis* in all samples

The human parasite *S. stercoralis* is often associated with low or inconsistent levels of nematode material or DNA in host faeces^91,92^. To assess if shotgun metagenomics is suitable for detecting *S. stercoralis*, we tested seven faecal samples collected from humans in Thailand or that had recently returned from Brunei with qPCR- or microscopy-confirmed *S. stercoralis* infections.

Only three samples yielded positive results with either Kraken2 (50RPM threshold), EukDetect, and/or mitochondrial mapping (**Fig. 3e-h**). For the Thailand samples (CKH130, CKH21, KKU01, LWH76), the two true-positive hits corresponded to the samples with the highest infection intensity (**Table 1**). Kraken2 generated putative false positive assignments to multiple species (FPR = 72%, **Fig. 3f**). Two out of seven samples assigned *S. stercoralis* correctly using all three of the analytical methods tested but four out of seven *S. stercoralis*-positive samples were not identified by shotgun metagenomics using any method. (**Fig. 3e-h**). Our results indicate that shotgun metagenomics is unreliable for accurate *S. stercoralis* detection based on the analytical methods we’ve used.

### 2.6 Genome assembly quality is important

Before applying Kraken2 to the human faecal samples, scaffolds <10kb were removed from the reference database because smaller scaffolds in genome assemblies often represent bacterial contamination^47^. To investigate if false positive assignments could be reduced further, we compared Kraken2 taxonomic assignment after removing scaffolds shorter than 10kb, 20kb and 40kb. The FPR of 14.2% (<10 kb) 15.3% (<20 kb) and 12.2% (40 kb) indicated that filtering of scaffolds up to 40 kb had little effect on the reducing FPR overall (**Additional File 10**).

Two samples falsely identified *P. trichosuri* (**Fig. 3b**) and false positive assignment of *P. trichosuri* was reduced by 24% (LWH76) and 38.7% (CKH130) when scaffolds <20kb were removed, and by 68.1% (LWH76) and 81.9% (CKH130), when scaffolds <40kb were removed. However, false positive assignment to *P. trichosuri* was still higher than alignments to *S. stercoralis*, indicating that removal of longer scaffolds did improve taxonomic assignment of *P. trichosuri*, but was insufficient for reliable STH detection (**Additional File 10**). The sample with the highest false assignment to *P. trichosuri* (LWH76) aligned 180,246 reads to 530 of the 1,810 scaffolds in the *P. trichosuri* genome. Of these, 524 scaffolds were <40 kb in length, and the longest scaffold was 1.04 Mb (**Supplementary Figure S3, Additional File 1, Additional File 10**). Overall, 93.4% of *P. trichosuri*-aligned reads were of bacterial origin with 91.22% mean sequence identity to *Brevundimonas* spp., which are ubiquitous in the environment (**Supplementary Figure S4, Additional File 1, Additional File 10**). These results indicate that false positive alignments occur due to contamination from bacterial sequences incorporated into eukaryotic genome assemblies.

## 4. Discussion

Shotgun metagenomics has potential for detecting STH infections and understanding microbiota-STH interactions^48,50–52,64–66^. Method validation is essential for accurate taxonomic assignment, maximising sensitivity and specificity for the application of shotgun metagenomics as a diagnostic tool. We have compared the analytical approach and sequencing technologies for shotgun metagenomics for two infection intensities in a laboratory model of STH infection using *S. ratti*, and tested human faecal samples with confirmed (using qPCR or formol-ether concentration) infections for at least one of four STHs.

Mitochondrial mapping of short-read metagenomics samples was the most accurate method of STH detection in rats infected with *S. ratti*, Other methods based on k-mer identification (Kraken2), marker genes (EukDetect) and protein sequences (DIAMOND+MEGAN) performed less well at a lower infection intensity. These results were mirrored in STH-infected human samples, where mitochondrial genome mapping was consistently the most effective method at minimising false positive and false negative detection in faecal samples infected with *Ascaris spp* and *N. americanus*. This supports previous work where mitochondrial mapping outperformed other methods of taxonomic assignment and likely reflects the high copy numbers of the mitochondrial genome present, thus increasing the chance of detection^47,93^.

The enhanced performance of mitochondrial mapping may also reflect the fact that mitochondrial genomes are often more complete and better curated than nuclear assemblies for many parasitic species. The mitochondrial mapping method is limited to detection of species with available mitochondrial genome sequences. Often, the mitochondrial and nuclear genomes of human and livestock parasites are more readily available, due to sampling bias of medically and economically important parasites. However, expansion of this method for less well-studied or rarer parasites could be limited by reference data availability. In 2024, there were complete mitochondrial genomes for 255 nematodes, 126 cestodes, 101 trematodes, and 24 acanthocephalan species available on NCBI, but up to 350,000 helminth species are estimated to exist^94^. Where mitochondrial genomes are not available, caution should be taken when interpreting taxonomic assignments, and method validation is recommended.

Nuclear genomes of helminths are frequently fragmented, contaminated with microbial sequences or incomplete, which can reduce the sensitivity and specificity of k-mer-, marker-, or protein-based detection methods. This discrepancy underscores the need for more complete and well-annotated nuclear genome assemblies if shotgun metagenomics is to be used as a reliable and generalisable detection strategy across diverse parasite taxa. False positive identification due to poor quality genome assemblies is an issue and poses a barrier to using shotgun metagenomics for detecting eukaryotes in primarily bacterial environments. Where eukaryote reference genomes contain bacterial contamination from common environmental sources, such as the *Brevundimonas* contamination in the *P. trichosuri* genome, signals for the true positive species present may be negligible. Bacterial contamination in eukaryotic genomes is increasingly documented^95,96^. While removing smaller scaffolds from reference databases does improve false positive detection rates, it does not resolve the issue. Continued efforts to refine and curate reference genomes will benefit shotgun metagenomics studies concerned with accurate identification of eukaryotes.

A remaining problem is that only a small proportion of the DNA, and subsequently the sequenced reads to be analysed in samples, are of parasite origin. In samples where DNA levels are expected to be low, detection was unreliable. Specifically, (i) in the *S. ratti* laboratory system, low infection intensity was difficult to detect using 3 out of 4 methods tested, (ii) *T. trichiura* could not be detected in human faecal samples - *T. trichiura* eggs are resilient compared to other STHs^97^, which can lead to difficulties in releasing DNA e.g. a previous attempt to detect *T. trichiura* using metagenomics obtained few reads^98^; and (iii) *S. stercoralis* was more readily detected in samples with higher larval recovery. Together these results support that the level of STH material, and thus DNA present in samples is important for shotgun-based STH detection. Future efforts focusing on enhancing recovery of parasite material before DNA extraction or during sequencing could resolve this. Development of laboratory based methods, not explored here, to concentrate parasite material and DNA recovery could further aide detection methods. Recent studies have investigated approaches to increase parasite-derived DNA signal in complex samples, notably through hybridization capture^99^ and refinement of metabarcoding and library-preparation workflows^100^. These enrichment and targeted sequencing techniques highlight promising avenues for improving parasite detection sensitivity in shotgun metagenomics. These methods will still require comprehensive validation.

Long read metagenomic sequencing approaches have potential to distinguish deeper taxonomic levels such as at a strain level, or could provide additional information about the diversity of parasite populations e.g. full ribosomal RNA coding regions^101^, which could aid detection of less well characterised parasites. Long-reads can also improve taxonomic recall^102^, but do not appear to eliminate false positive assignments. However, the ONT metagenomics used here did not reliably detect *S. ratti* true-positives. Hybrid approaches, which combine short and long read data successfully detected true-positives and no false positives, but this was mostly attributed to the short-reads present, thus offering no benefit of long-read sequencing. Long-read data analysis for eukaryotic metagenomics requires further development to unlock any potential benefits. However, currently the increased computational complexity and analysis time, high cost, higher input levels and generation of low read numbers further impacts feasibility for large-scale or resource-limited studies with long-read metagenomics, compared to short read ,metagenomic. The development of more accurate and computationally efficient tools tailored to long-read eukaryotic metagenomic data should be a priority to fully realise the potential of long-read and hybrid sequencing for STH detection.

## Authors’ contributions

KOB, AE, JY and VLH conceived and designed the project; KOB, WN, YL, MV, CB, BS, PSN collected samples and carried out laboratory work; KOB & KR carried out long read sequencing; KOB, AE, MD, GK, KR, MW carried out analysis, KOB, AE, RW, VH wrote the manuscript.

## Supporting Information Captions

### Additional file 1 - Supplementary methods , figures and tables

Supplementary Methods

Supplementary Figure S1 – S2: Alpha and beta diversity results for short-read metagenomics of rat faecal samples

Supplementary Figure S3: Coverage of LWH76 alignments to the *P. trichosuri* Genome show that the Majority of Hits are to Small Scaffolds

Supplementary Figure 4: Alignment of *P. trichosuri* Contigs to the *B. naejangsanensis* Genome Indicates Bacterial Contamination

Supplementary Tables S1-3: Short-read metagenomics of rat faecal samples

Supplementary Table S4: Short-read metagenomic shotgun reads sequenced from human samples with positive PCR for *Strongyloides stercoralis* infection

Supplementary Table S5: Long-read metagenomics of rat faecal samples - raw sequencing data and host removed/filtered sequence data

Supplementary Table S6: Parasite genomes included in custom Kraken2 database Supplementary Table S7: Hybrid Assembly Metrics

Supplementary Table S8: Short-read metagenomic shotgun reads

Supplementary Table S9: Larval counts for S. ratti infectiosn at 6 and 11 days post-infection at standard and low infection intensities

**Additional file 2 -** Coverage values and reads aligned per million reads from Kraken2, DIAMOND+MEGAN. EukDetect and mitochondrial mapping using short-read metagenomics

**Additional file 3** - Taxonomic assignments for long read data reported as Bases per million and normalised coverage values.

**Additional file 4 -** Taxonomic assignment of binned sequences from MetaBAT2

**Additional file 5 -** BLAST results of the unbinned sequences

**Additional file 6 -** Measures of sensitivity, specificity, precision, false positive detection rate (FPR), and F1 scores for short-read analysis

**Additional file 7 -** Measures of sensitivity, specificity, precision, false positive detection rate (FPR), and F1 scores long-read and long-short read hybrid analysis. **Additional file 8 -** Coverage values and reads aligned per million reads from Kraken2, DIAMOND+MEGAN. EukDetect and mitochondrial mapping using ,etagenomic shotgun human samples.

**Additional file 9 -** Measures of sensitivity, specificity, precision, false positive detection rate (FPR), and F1 scores from human samples.

**Additional file 10 -** Kraken2 taxonomic assignment after removing scaffolds shorter than 10kb, 20kb and 40kb. The FPR of 14.2% (<10 kb) 15.3% (<20 kb) and 12.2% (40 kb)

## Notes

### Competing Interest Statement

The authors have declared no competing interest.

## References

1. Montresor A, Mupfasoni D, Mikhailov A, et al. The global progress of soil-transmitted helminthiases control in 2020 and World Health Organization targets for 2030. Babu S, ed. PLoS Negl Trop Dis. 2020;14(8):e0008505. doi:10.1371/journal.pntd.0008505

2. Soares FA, Benitez ADN, Santos BMD, et al. A historical review of the techniques of recovery of parasites for their detection in human stools. Rev Soc Bras Med Trop. 2020;53:e20190535. doi:10.1590/0037-8682-0535-2019

3. Gonçalves AQ, Abellana R, Pereira-da-Silva HD, et al. Comparison of the performance of two spontaneous sedimentation techniques for the diagnosis of human intestinal parasites in the absence of a gold standard. Acta Tropica. 2014;131:63–70. doi:10.1016/j.actatropica.2013.11.026

4. Speich B, Ali SM, Ame SM, Albonico M, Utzinger J, Keiser J. Quality control in the diagnosis of Trichuris trichiura and Ascaris lumbricoides using the Kato-Katz technique: experience from three randomised controlled trials. Parasites Vectors. 2015;8(1):82. doi:10.1186/s13071-015-0702-z

5. Momčilović S, Cantacessi C, Arsić-Arsenijević V, Otranto D, Tasić-Otašević S. Rapid diagnosis of parasitic diseases: current scenario and future needs. Clinical Microbiology and Infection. 2019;25(3):290–309. doi:10.1016/j.cmi.2018.04.028

6. Lim MD, Brooker SJ, Belizario VY, et al. Diagnostic tools for soil-transmitted helminths control and elimination programs: A pathway for diagnostic product development. Maruyama H, ed. PLoS Negl Trop Dis. 2018;12(3):e0006213. doi:10.1371/journal.pntd.0006213

7. Siddiqui AA, Berk SL. Diagnosis of *Strongyloides stercoralis* Infection. CLIN INFECT DIS. 2001;33(7):1040–1047. doi:10.1086/322707

8. Nikolay B, Brooker SJ, Pullan RL. Sensitivity of diagnostic tests for human soil-transmitted helminth infections: a meta-analysis in the absence of a true gold standard. International Journal for Parasitology. 2014;44(11):765–774. doi:10.1016/j.ijpara.2014.05.009

9. Buonfrate D, Bisanzio D, Giorli G, et al. The Global Prevalence of Strongyloides stercoralis Infection. Pathogens. 2020;9(6):468. doi:10.3390/pathogens9060468

10. Bisoffi Z, Buonfrate D, Montresor A, et al. Strongyloides stercoralis: A Plea for Action. Lammie PJ, ed. PLoS Negl Trop Dis. 2013;7(5):e2214. doi:10.1371/journal.pntd.0002214

11. Bush A, Compson ZG, Monk WA, et al. Studying Ecosystems With DNA Metabarcoding: Lessons From Biomonitoring of Aquatic Macroinvertebrates. Front Ecol Evol. 2019;7:434. doi:10.3389/fevo.2019.00434

12. Buchan BW, Ledeboer NA. Emerging Technologies for the Clinical Microbiology Laboratory. Clin Microbiol Rev. 2014;27(4):783–822. doi:10.1128/CMR.00003-14

13. Rashwan N, Diawara A, Scott ME, Prichard RK. Isothermal diagnostic assays for the detection of soil-transmitted helminths based on the SmartAmp2 method. Parasites Vectors. 2017;10(1):496. doi:10.1186/s13071-017-2420-1

14. O’Connell EM, Nutman TB. Molecular Diagnostics for Soil-Transmitted Helminths. The American Society of Tropical Medicine and Hygiene. 2016;95(3):508–513. doi:10.4269/ajtmh.16-0266

15. Ji Y, Ashton L, Pedley SM, et al. Reliable, verifiable and efficient monitoring of biodiversity via metabarcoding. Holyoak M, ed. Ecology Letters. 2013;16(10):1245–1257. doi:10.1111/ele.12162

16. Fernández-Soto P, Fernández-Medina C, Cruz-Fernández S, et al. Whip-LAMP: a novel LAMP assay for the detection of Trichuris muris-derived DNA in stool and urine samples in a murine experimental infection model. Parasites Vectors. 2020;13(1):552. doi:10.1186/s13071-020-04435-1

17. Aird D, Ross MG, Chen WS, et al. Analyzing and minimizing PCR amplification bias in Illumina sequencing libraries. Genome Biol. 2011;12(2):R18. doi:10.1186/gb-2011-12-2-r18

18. Parada AE, Needham DM, Fuhrman JA. Every base matters: assessing small subunit RRNA primers for marine microbiomes with mock communities, time series and global field samples. Environmental Microbiology. 2016;18(5):1403–1414. doi:10.1111/1462-2920.13023

19. Kounosu A, Murase K, Yoshida A, Maruyama H, Kikuchi T. Improved 18S and 28S rDNA primer sets for NGS-based parasite detection. Sci Rep. 2019;9(1):15789. doi:10.1038/s41598-019-52422-z

20. Konstantinidis KT, Tiedje JM. Prokaryotic taxonomy and phylogeny in the genomic era: advancements and challenges ahead. Current Opinion in Microbiology. 2007;10(5):504–509. doi:10.1016/j.mib.2007.08.006

21. Boers SA, Jansen R, Hays JP. Understanding and overcoming the pitfalls and biases of next-generation sequencing (NGS) methods for use in the routine clinical microbiological diagnostic laboratory. Eur J Clin Microbiol Infect Dis. 2019;38(6):1059–1070. doi:10.1007/s10096-019-03520-3

22. Raso G. Multiple parasite infections and their relationship to self-reported morbidity in a community of rural Cote d’Ivoire. International Journal of Epidemiology. 2004;33(5):1092–1102. doi:10.1093/ije/dyh241

23. Finkelman FD, Shea-Donohue T, Morris SC, et al. Interleukin-4- and interleukin-13-mediated host protection against intestinal nematode parasites. Immunological Reviews. 2004;201(1):139–155. doi:10.1111/j.0105-2896.2004.00192.x

24. Vacca F, Le Gros G. Tissue-specific immunity in helminth infections. Mucosal Immunology. 2022;15(6):1212–1223. doi:10.1038/s41385-022-00531-w

25. Quince C, Walker AW, Simpson JT, Loman NJ, Segata N. Shotgun metagenomics, from sampling to analysis. Nat Biotechnol. 2017;35(9):833–844. doi:10.1038/nbt.3935

26. Riesenfeld CS, Schloss PD, Handelsman J. Metagenomics: Genomic Analysis of Microbial Communities. Annu Rev Genet. 2004;38(1):525–552. doi:10.1146/annurev.genet.38.072902.091216

27. Colston TJ, Jackson CR. Microbiome evolution along divergent branches of the vertebrate tree of life: what is known and unknown. Molecular Ecology. 2016;25(16):3776–3800. doi:10.1111/mec.13730

28. Jovel J, Patterson J, Wang W, et al. Characterization of the Gut Microbiome Using 16S or Shotgun Metagenomics. Front Microbiol. 2016;7. doi:10.3389/fmicb.2016.00459

29. Mendes LW, Braga LPP, Navarrete AA, Souza DGD, Silva GGZ, Tsai SM. Using Metagenomics to Connect Microbial Community Biodiversity and Functions. Current Issues in Molecular Biology. 2017:103–118. doi:10.21775/cimb.024.103

30. Mongkolrattanothai K, Naccache SN, Bender JM, et al. Neurobrucellosis: Unexpected Answer From Metagenomic Next-Generation Sequencing. JPIDSJ. January 2017:piw066. doi:10.1093/jpids/piw066

31. Sczyrba A, Hofmann P, Belmann P, et al. Critical Assessment of Metagenome Interpretation—a benchmark of metagenomics software. Nat Methods. 2017;14(11):1063–1071. doi:10.1038/nmeth.4458

32. Meyer F, Fritz A, Deng ZL, et al. Critical Assessment of Metagenome Interpretation: the second round of challenges. Nat Methods. 2022;19(4):429–440. doi:10.1038/s41592-022-01431-4

33. Lu J, Rincon N, Wood DE, et al. Metagenome analysis using the Kraken software suite. Nat Protoc. 2022;17(12):2815–2839. doi:10.1038/s41596-022-00738-y

34. Keegan KP, Glass EM, Meyer F. MG-RAST, a Metagenomics Service for Analysis of Microbial Community Structure and Function. In: Martin F, Uroz S, eds. Microbial Environmental Genomics (MEG). Vol 1399. Methods in Molecular Biology. New York, NY: Springer New York; 2016:207–233. doi:10.1007/978-1-4939-3369-3_13

35. Beghini F, McIver LJ, Blanco-Míguez A, et al. Integrating taxonomic, functional, and strain-level profiling of diverse microbial communities with bioBakery 3. eLife. 2021;10:e65088. doi:10.7554/eLife.65088

36. Pérez-Cobas AE, Gomez-Valero L, Buchrieser C. Metagenomic approaches in microbial ecology: an update on whole-genome and marker gene sequencing analyses. Microbial Genomics. 2020;6(8). doi:10.1099/mgen.0.000409

37. Grützke J, Malorny B, Hammerl JA, et al. Fishing in the Soup – Pathogen Detection in Food Safety Using Metabarcoding and Metagenomic Sequencing. Front Microbiol. 2019;10:1805. doi:10.3389/fmicb.2019.01805

38. Liu D, Zhou H, Xu T, et al. Multicenter assessment of shotgun metagenomics for pathogen detection. eBioMedicine. 2021;74:103649. doi:10.1016/j.ebiom.2021.103649

39. Yen S, Johnson JS. Metagenomics: a path to understanding the gut microbiome. Mamm Genome. 2021;32(4):282–296. doi:10.1007/s00335-021-09889-x

40. Laforest-Lapointe I, Arrieta MC. Microbial Eukaryotes: a Missing Link in Gut Microbiome Studies. mSystems. 2018;3(2):e00201–17. doi:10.1128/mSystems.00201-17

41. Reynoso-García J, Santiago-Rodriguez TM, Narganes-Storde Y, Cano RJ, Toranzos GA. Edible flora in pre-Columbian Caribbean coprolites: Expected and unexpected data. Hart JP, ed. PLoS ONE. 2023;18(10):e0292077. doi:10.1371/journal.pone.0292077

42. Mann AE, Fellows Yates JA, Fagernäs Z, Austin RM, Nelson EA, Hofman CA. Do I have something in my teeth? The trouble with genetic analyses of diet from archaeological dental calculus. Quaternary International. 2023;653-654:33–46. doi:10.1016/j.quaint.2020.11.019

43. Srivathsan A, Ang A, Vogler AP, Meier R. Fecal metagenomics for the simultaneous assessment of diet, parasites, and population genetics of an understudied primate. Front Zool. 2016;13(1):17. doi:10.1186/s12983-016-0150-4

44. Serite CP, Emami-Khoyi A, Ntshudisane OK, et al. eDNA metabarcoding vs metagenomics: an assessment of dietary competition in two estuarine pipefishes. Front Mar Sci. 2023;10:1116741. doi:10.3389/fmars.2023.1116741

45. Chua PYS, Crampton-Platt A, Lammers Y, Alsos IG, Boessenkool S, Bohmann K. Metagenomics: A viable tool for reconstructing herbivore diet. Molecular Ecology Resources. 2021;21(7):2249–2263. doi:10.1111/1755-0998.13425

46. Wylezich C, Caccio SM, Walochnik J, Beer M, Höper D. Untargeted metagenomics shows a reliable performance for synchronous detection of parasites. Parasitol Res. 2020;119(8):2623–2629. doi:10.1007/s00436-020-06754-9

47. Papaiakovou M, Fraija-Fernández N, James K, et al. Evaluation of genome skimming to detect and characterise human and livestock helminths. International Journal for Parasitology. 2023;53(2):69–79. doi:10.1016/j.ijpara.2022.12.002

48. Lind AL, Pollard KS. Accurate and sensitive detection of microbial eukaryotes from whole metagenome shotgun sequencing. Microbiome. 2021;9(1):58. doi:10.1186/s40168-021-01015-y

49. Franssen FFJ, Janse I, Janssen D, et al. Mining Public Metagenomes for Environmental Surveillance of Parasites: A Proof of Principle. Front Microbiol. 2021;12:622356. doi:10.3389/fmicb.2021.622356

50. Mthethwa NP, Amoah ID, Reddy P, Bux F, Kumari S. A review on application of next-generation sequencing methods for profiling of protozoan parasites in water: Current methodologies, challenges, and perspectives. Journal of Microbiological Methods. 2021;187:106269. doi:10.1016/j.mimet.2021.106269

51. Wylezich C, Belka A, Hanke D, Beer M, Blome S, Höper D. Metagenomics for broad and improved parasite detection: a proof-of-concept study using swine faecal samples. International Journal for Parasitology. 2019;49(10):769–777. doi:10.1016/j.ijpara.2019.04.007

52. MetaHIT Consortium, Qin J, Li R, et al. A human gut microbial gene catalogue established by metagenomic sequencing. Nature. 2010;464(7285):59–65. doi:10.1038/nature08821

53. Steinegger M, Salzberg SL. Terminating contamination: large-scale search identifies more than 2,000,000 contaminated entries in GenBank. Genome Biol. 2020;21(1):115. doi:10.1186/s13059-020-02023-1

54. Bálint B, Merényi Z, Hegedüs B, et al. ContScout: sensitive detection and removal of contamination from annotated genomes. Nat Commun. 2024;15(1):936. doi:10.1038/s41467-024-45024-5

55. Lu J, Salzberg SL. Removing contaminants from databases of draft genomes. Sun F, ed. PLoS Comput Biol. 2018;14(6):e1006277. doi:10.1371/journal.pcbi.1006277

56. Mikheyev AS, Tin MMY. A first look at the Oxford Nanopore MinION sequencer. Molecular Ecology Resources. 2014;14(6):1097–1102. doi:10.1111/1755-0998.12324

57. Rhoads A, Au KF. PacBio Sequencing and its Applications. Genomics, Proteomics & Bioinformatics. 2015;13(5):278–289. doi:10.1016/j.gpb.2015.08.002

58. Hu T, Chitnis N, Monos D, Dinh A. Next-generation sequencing technologies: An overview. Human Immunology. 2021;82(11):801–811. doi:10.1016/j.humimm.2021.02.012

59. Marais G, Hardie D, Brink A. A case for investment in clinical metagenomics in low-income and middle-income countries. The Lancet Microbe. 2023;4(3):e192–e199. doi:10.1016/S2666-5247(22)00328-7

60. Viney M, Kikuchi T. *Strongyloides ratti* and *S. venezuelensis* – rodent models of *Strongyloides* infection. Parasitology. 2017;144(3):285–294. doi:10.1017/S0031182016000020

61. Hunt VL, Tsai IJ, Coghlan A, et al. The genomic basis of parasitism in the Strongyloides clade of nematodes. Nat Genet. 2016;48(3):299–307. doi:10.1038/ng.3495

62. Viney ME, Matthews BE, Walliker D. On the biological and biochemical nature of cloned populations of *Strongyloides ratti*. J Helminthol. 1992;66(1):45–52. doi:10.1017/S0022149X00012554

63. Langmead B, Salzberg SL. Fast gapped-read alignment with Bowtie 2. Nat Methods. 2012;9(4):357–359. doi:10.1038/nmeth.1923

64. Li H, Handsaker B, Wysoker A, et al. The Sequence Alignment/Map format and SAMtools. Bioinformatics. 2009;25(16):2078–2079. doi:10.1093/bioinformatics/btp352

65. Quinlan AR, Hall IM. BEDTools: a flexible suite of utilities for comparing genomic features. Bioinformatics. 2010;26(6):841–842. doi:10.1093/bioinformatics/btq033

66. Lanfear R, Schalamun M, Kainer D, Wang W, Schwessinger B. MinIONQC: fast and simple quality control for MinION sequencing data. Hancock J, ed. Bioinformatics. 2019;35(3):523–525. doi:10.1093/bioinformatics/bty654

67. Andrews, S. FastQC: a quality control tool for high throughput sequence data. http://www.bioinformatics.babraham.ac.uk/projects/fastqc.

68. De Coster W, D’Hert S, Schultz DT, Cruts M, Van Broeckhoven C. NanoPack: visualizing and processing long-read sequencing data. Berger B, ed. Bioinformatics. 2018;34(15):2666–2669. doi:10.1093/bioinformatics/bty149

69. Li H. Minimap2: pairwise alignment for nucleotide sequences. Birol I, ed. Bioinformatics. 2018;34(18):3094–3100. doi:10.1093/bioinformatics/bty191

70. Wood DE, Lu J, Langmead B. Improved metagenomic analysis with Kraken 2. Genome Biol. 2019;20(1):257. doi:10.1186/s13059-019-1891-0

71. Bağcı C, Patz S, Huson DH. DIAMOND+MEGAN: Fast and Easy Taxonomic and Functional Analysis of Short and Long Microbiome Sequences. Current Protocols. 2021;1(3):e59. doi:10.1002/cpz1.59

72. Lind AL, Pollard KS. Accurate and sensitive detection of microbial eukaryotes from whole metagenome shotgun sequencing. Microbiome. 2021;9(1):58. doi:10.1186/s40168-021-01015-y

73. Howe KL, Bolt BJ, Shafie M, Kersey P, Berriman M. WormBase ParaSite − a comprehensive resource for helminth genomics. Molecular and Biochemical Parasitology. 2017;215:2–10. doi:10.1016/j.molbiopara.2016.11.005

74. McMurdie PJ, Holmes S. phyloseq: An R Package for Reproducible Interactive Analysis and Graphics of Microbiome Census Data. Watson M, ed. PLoS ONE. 2013;8(4):e61217. doi:10.1371/journal.pone.0061217

75. Winter DJ. rentrez: An R package for the NCBI eUtils API. August 2017. doi:10.7287/peerj.preprints.3179v1

76. Oksanen J, Simpson G, Blanchet F, et al. vegan: Community Ecology Package. R package version 2.8-0. https://vegandevs.github.io/vegan/.

77. Wickham H. ggplot2: Elegant Graphics for Data Analysis. https://ggplot2.tidyverse.org.

78. Buchfink B, Xie C, Huson DH. Fast and sensitive protein alignment using DIAMOND. Nat Methods. 2015;12(1):59–60. doi:10.1038/nmeth.3176

79. Huson DH, Beier S, Flade I, et al. MEGAN Community Edition - Interactive Exploration and Analysis of Large-Scale Microbiome Sequencing Data. Poisot T, ed. PLoS Comput Biol. 2016;12(6):e1004957. doi:10.1371/journal.pcbi.1004957

80. Lind A. EukDetect database. 2021:0 Bytes. doi:10.6084/M9.FIGSHARE.13626812

81. Li H. Aligning sequence reads, clone sequences and assembly contigs with BWA-MEM. 2013. doi:10.48550/ARXIV.1303.3997

82. Simmonds SE, Ly L, Beaulaurier J, et al. CZ ID: a cloud-based, no-code platform enabling advanced long read metagenomic analysis. March 2024. doi:10.1101/2024.02.29.579666

83. Kalantar KL, Carvalho T, De Bourcy CFA, et al. IDseq—An open source cloud-based pipeline and analysis service for metagenomic pathogen detection and monitoring. GigaScience. 2020;9(10):giaa111. doi:10.1093/gigascience/giaa111

84. Nurk S, Meleshko D, Korobeynikov A, Pevzner PA. metaSPAdes: a new versatile metagenomic assembler. Genome Res. 2017;27(5):824–834. doi:10.1101/gr.213959.116

85. Mikheenko A, Saveliev V, Gurevich A. MetaQUAST: evaluation of metagenome assemblies. Bioinformatics. 2016;32(7):1088–1090. doi:10.1093/bioinformatics/btv697

86. Kang DD, Li F, Kirton E, et al. MetaBAT 2: an adaptive binning algorithm for robust and efficient genome reconstruction from metagenome assemblies. PeerJ. 2019;7:e7359. doi:10.7717/peerj.7359

87. Parks DH, Imelfort M, Skennerton CT, Hugenholtz P, Tyson GW. CheckM: assessing the quality of microbial genomes recovered from isolates, single cells, and metagenomes. Genome Res. 2015;25(7):1043–1055. doi:10.1101/gr.186072.114

88. Rosa BA, Supali T, Gankpala L, et al. Differential human gut microbiome assemblages during soil-transmitted helminth infections in Indonesia and Liberia. Microbiome. 2018;6(1):33. doi:10.1186/s40168-018-0416-5

89. Bolger AM, Lohse M, Usadel B. Trimmomatic: a flexible trimmer for Illumina sequence data. Bioinformatics. 2014;30(15):2114–2120. doi:10.1093/bioinformatics/btu170

90. Doyle SR, Søe MJ, Nejsum P, et al. Population genomics of ancient and modern Trichuris trichiura. Nat Commun. 2022;13(1):3888. doi:10.1038/s41467-022-31487-x

91. Dreyer G, Fernandes-Silva E, Alves S, Rocha A, Albuquerque R, Addiss D. Patterns of detection of Strongyloides stercoralis in stool specimens: implications for diagnosis and clinical trials. J Clin Microbiol. 1996;34(10):2569–2571. doi:10.1128/jcm.34.10.2569-2571.1996

92. Uparanukraw P, Phongsri S, Morakote N. Fluctuations of larval excretion in Strongyloides stercoralis infection. Am J Trop Med Hyg. 1999;60(6):967–973. doi:10.4269/ajtmh.1999.60.967

93. Doyle SR, Søe MJ, Nejsum P, et al. Population genomics of ancient and modern Trichuris trichiura. Nat Commun. 2022;13(1):3888. doi:10.1038/s41467-022-31487-x

94. Carlson CJ, Dallas TA, Alexander LW, Phelan AL, Phillips AJ. What would it take to describe the global diversity of parasites? Proc R Soc B. 2020;287(1939):20201841. doi:10.1098/rspb.2020.1841

95. Percudani R. A Microbial Metagenome ( *Leucobacter* sp.) in *Caenorhabditis* Whole Genome Sequences. Bioinform Biol Insights. 2013;7:BBI.S11064. doi:10.4137/BBI.S11064

96. Artamonova II, Mushegian AR. Genome Sequence Analysis Indicates that the Model Eukaryote Nematostella vectensis Harbors Bacterial Consorts. Appl Environ Microbiol. 2013;79(22):6868–6873. doi:10.1128/AEM.01635-13

97. Kaisar MMM, Brienen EAT, Djuardi Y, et al. Improved diagnosis of *Trichuris trichiura* by using a bead-beating procedure on ethanol preserved stool samples prior to DNA isolation and the performance of multiplex real-time PCR for intestinal parasites. Parasitology. 2017;144(7):965–974. doi:10.1017/S0031182017000129

98. Chessa D, Murgia M, Sias E, et al. Metagenomics and microscope revealed T. trichiura and other intestinal parasites in a cesspit of an Italian nineteenth century aristocratic palace. Sci Rep. 2020;10(1):12656. doi:10.1038/s41598-020-69497-8

99. Papaiakovou M, Waeschenbach A, Anderson RM, et al. Enrichment of Helminth Mitochondrial Genomes From Faecal Samples Using Hybridisation Capture. Molecular Ecology Resources. 2025;25(8):e70005. doi:10.1111/1755-0998.70005

100. Kang D, Choi JH, Kim M, et al. Optimization of 18 S rRNA metabarcoding for the simultaneous diagnosis of intestinal parasites. Sci Rep. 2024;14(1):25049. doi:10.1038/s41598-024-76304-1

101. Gaonkar CC, Campbell L. A full-length 18S ribosomal DNA metabarcoding approach for determining protist community diversity using Nanopore sequencing. Ecology and Evolution. 2024;14(4):e11232. doi:10.1002/ece3.11232

102. Pearman WS, Freed NE, Silander OK. Testing the advantages and disadvantages of short- and long- read eukaryotic metagenomics using simulated reads. BMC Bioinformatics. 2020;21(1):220. doi:10.1186/s12859-020-3528-4

